# Nonequilibrium brain dynamics elicited as the origin of perturbative complexity

**DOI:** 10.1101/2024.11.29.625885

**Authors:** Wiep Stikvoort, Eider Pérez-Ordoyo, Iván Mindlin, Anira Escrichs, Jacobo D. Sitt, Morten L. Kringelbach, Gustavo Deco, Yonatan Sanz Perl

**Affiliations:** Center for Brain and Cognition, Computational Neuroscience Group, Universitat Pompeu Fabra, Barcelona, Spain; Paris Brain Institute, ICM, Inserm, CNRS, Sorbonne Université, Paris, France; Department of Psychiatry, University of Oxford, Oxford, UK; Centre for Eudoimonia and Human Flourishing, Linacre College, University of Oxford, Oxford, UK; Centre for Music in the Brain, Aarhus University, Aarhus, Denmark; Institució Catalana de la Recerca i Estudis Avançats (ICREA), Barcelona, Spain; Buenos Aires Physics Institute and Physics Department, University of Buenos Aires, Buenos Aires, Argentina; Universidad de San Andres, Buenos Aires, Argentina; Institut de Cerveau et de la Moelle epiniere, ICM, Paris, France

**Keywords:** *whole-brain modelling*, *effective connectivity*, *asymmetry*, *PCI*, *nonequilibrium*, *consciousness*

## Abstract

Assessing the level of consciousness someone is in, is not a trivial question and physicians have to rely on behavioural evaluations instead of quantifiable metrics. Many studies have empirically investigated measures related to the complexity elicited after the brain is stimulated to quantify and assess the level of consciousness across different states. Here we hypothesized that the level of non-equilibrium dynamics of the unperturbed brain already contains the information needed to know how the system will react to an external stimulus. We created personalized whole-brain models fitted to resting state fMRI data recorded in participants in different states of reduced consciousness (such as deep sleep and disorders of consciousness) to infer the effective connections underlying their brain dynamics. We then measured the out-of-equilibrium nature of the unperturbed brain by evaluating the level of asymmetry of the inferred connectivity, the time irreversibility in each model and compared this with the elicited complexity generated after *in silico* perturbations. Crucially, we found that states of reduced consciousness had a lower level of asymmetry in their effective connectivities compared to control subjects, as well as a lower level of irreversibility in their simulated dynamics, and a lower complexity. We demonstrated that the asymmetry in the underlying connections drives the nonequilibrium state of the system and in turn the differences in complexity as a response to the external stimuli.

## Introduction

There is an ongoing debate in neuroscience about the connection between consciousness and brain complexity (1). In this case complexity is defined as the combined presence of integration and segregation and can be measured by computing the spatiotemporal complexity of brain activity (2) and emerges from the underlying critical dynamics (3). An established approach to investigate this relationship was proposed by Massimini and colleagues, who assessed the perturbation-elicited changes in global brain activity during different states of consciousness, such as wakefulness, sleep, anaesthesia, and disorders of consciousness (DoC) (4–6). Specifically, they computed the perturbational complexity index (PCI), which captures the significant differences in brain-wide spatiotemporal propagation of external stimulation, and they proposed it as an index of consciousness (4). Importantly, the PCI had high accuracy in distinguishing different levels of consciousness and the results were replicated in multiple studies, showing its sensitivity and usefulness (7–10). Although the PCI has been proven to be valuable and accurate, it requires an extensive clinical setup which allows for perturbation of the brain with Transcranial Magnetic Stimulation (TMS) while recording brain activity with electroencephalography (EEG).

The concepts of detailed balance and irreversibility from thermodynamics have been applied to assess a system’s ability to exhibit dynamic fluctuations between different states. When a system is said to violate detailed balance, it exhibits net fluxes of e.g. activity, between its configurations, thereby establishing an arrow of time and rendering the system irreversible (11). Studies of spontaneous brain activity in healthy subjects have demonstrated that the brain operates in such a nonequilibrium state (12).

The main goal of the present work is to establish whether PCI, as a marker of consciousness in the human brain, can be associated with the dynamical nature of brain activity before being perturbed. We hypothesize that the level of nonequilibrium dynamics of the unperturbed system determines the response to an external stimulus and in turn the elicited complexity (PCI). To this end, we leverage recent works that proposed different theoretically-grounded approaches to measure the level of nonequilibrium at the whole-brain scale, such as entropy production (13), violation of the Fluctuation Dissipation Theorem (FDT)(14) and temporal asymmetry of brain signals (11,13,14). In particular, FDT provides a direct link between the state of the system and its response to a stimulus following the Onsager derivation (15–17), equalizing the spontaneous fluctuations and the external perturbation (14). Based on these thermodynamic concepts, previous works have empirically demonstrated that time asymmetry is reduced during deep sleep (18) and in DoC patients (19). Here we focus on the mechanism behind this nonequilibrium nature of brain dynamics (20) and how this property can be related to PCI.

In this study, we used a whole-brain modelling approach, as opposed to the previously used empirical TMS-EEG setup. We created personalized whole-brain models fitted to functional Magnetic Resonance Imaging (fMRI) data, which allowed us to exhaustively investigate the perturbative *in silico* response of different states of consciousness. Crucially, we introduced asymmetry in the models creating time hierarchy organization and breaking the detailed balance, causing there to be a state of nonequilibrium (20). By incorporating the time asymmetry of brain dynamics, which reveal the generative underlying mechanism, asymmetry is introduced in the connections, named generative effective connectivity (gEC) (20). The gEC is responsible for balance of the causal interaction between brain regions and in turn the level of nonequilibrium in different states of consciousness.

Taking advantage of the thermodynamics framework as a natural way to quantify nonequilibrium of brain activity and its underlying mechanisms (18), we use the asymmetry of the gEC to explore its relationship with nonequilibrium dynamics and *in silico* PCI estimates at subject-level for participants in various states of reduced consciousness. Crucially, we found that lower levels of consciousness generally have a lower level of asymmetry in their generative fitted connectivities and are thus inherently closer to an equilibrium state in terms of thermodynamics. These models fitted to participants residing in a lower level of consciousness also have lower PCI values, as well as lower levels of nonequilibrium. Through our models we demonstrated that the asymmetry in the generative underlying connections is causing the emergent nonequilibrium dynamics and, at the same time, the differences in the complexity elicited by perturbations measured with PCI.

## Results

### Overview

We used two fMRI datasets with subjects residing in different states of consciousness: a Disorders of Consciousness (DoC) dataset containing data from subjects in a Minimally Conscious State (MCS) (N = 11), an Unresponsive Wakefulness State (UWS) (N = 10), or a control state (CNT) (N = 13), and a sleep dataset containing data from subjects residing in wakefulness (W) and deep sleep (N3) (N = 18). In both datasets the activity was parcellated into 90 Regions of Interest (ROI’s), using the AAL parcellation (21). A schematic overview of the pipeline can be found in Figure 1. The empirical fMRI data is used to calculate the time-lagged covariance, depicted in Figure 1A and 1B, this time-lag ensures that we have a causal inference in the infrastructure of our models. In other words, we included the time-shifted correlations between ROI’s from the empirical data allowing the model to have bidirectional connections. Activity is then simulated by representing each ROI as an oscillator, coupled to other oscillators through a connectivity matrix (see Methods).

**Figure 1.**
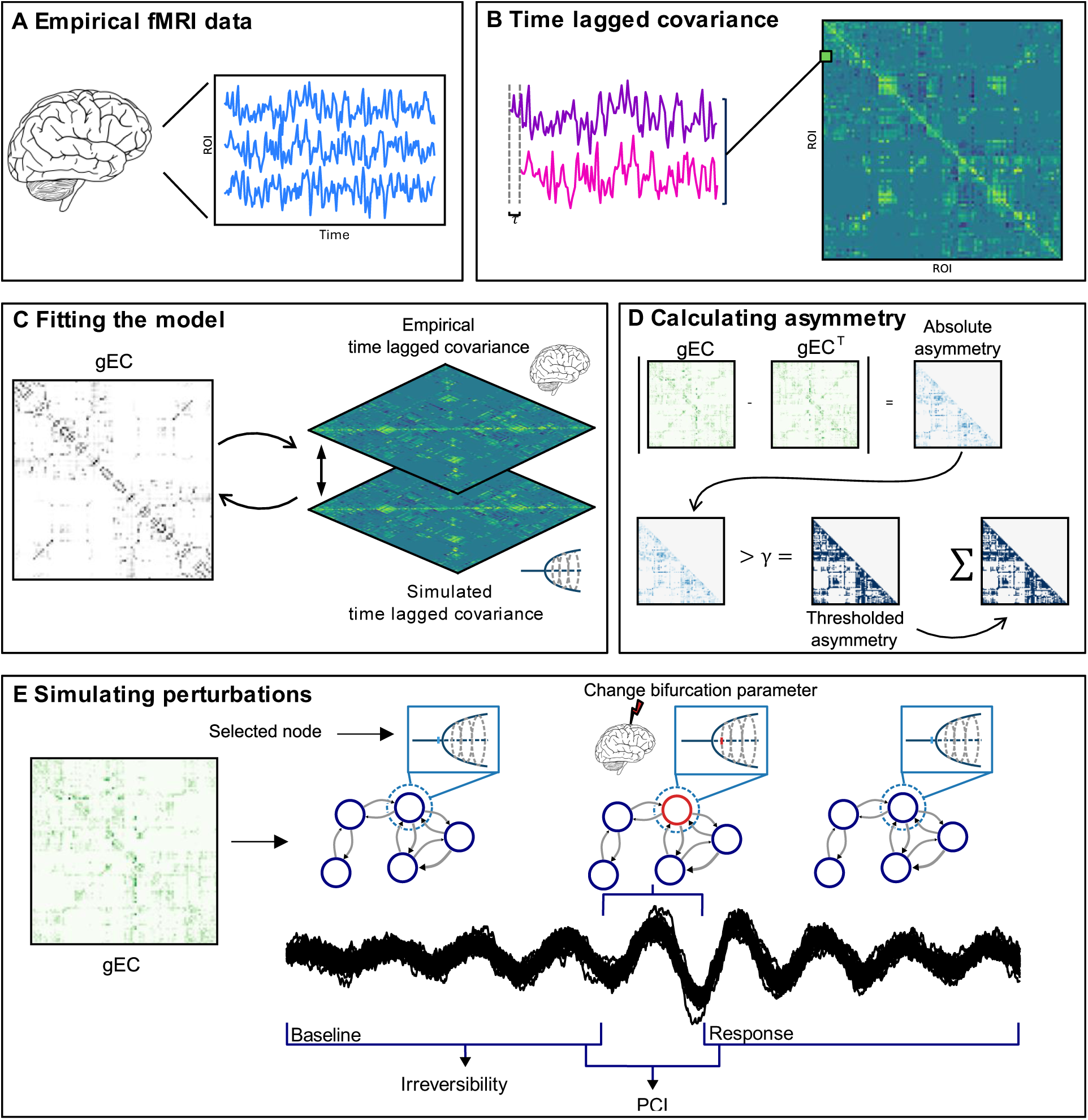
A schematic overview of the pipeline. A) fMRI data empirically retrieved from subjects in various states of consciousness, namely control (CNT), Minimally Conscious State (MCS), Unresponsive Wakefulness State (UWS), Wakefulness (W) and deep sleep (N3). B, C) To fit the models, the time lagged covariance is calculated from the empirical data and the simulated data. Through an iterative process in which the generative Effective Connectivity (gEC) is updated, the simulated time lagged covariance will resemble the empirical time lagged covariance as close as possible. D) The asymmetry of the gEC is calculated by thresholding the absolute values of the transposed gEC subtracted from its original form, the number of pairs of nodes that exceeded the threshold are summed which gives us the level of asymmetry in the model. E) The gEC is taken as the connectivity matrix in the Hopf model (C in eq. 2 and 3), in which each ROI is represented as a Stuart-Landau oscillator in the system. The model is perturbed by changing the value of the bifurcation parameter to a positive value for one of the nodes (a in eq. 2 and 3). This perturbation will transcend through the network, and by doing so we get a preperturbational and postperturbational time series, a baseline and a response window respectively, from which we can calculate the State Transition PCI. The irreversibility is calculated from the preperturbational time series, the resting state dynamics of the network. The differences in the gEC for each model will cause the irreversibility and PCI to be different.

Briefly, the fitting procedure starts with the Structural Connectivity (SC), which in this case is an averaged SC from a healthy group of subjects, and then we include the time-lagged covariance of all connections and update the weights using the values we retrieved from the empirical data, as shown in Figure 1C. Using the updated weights, the process is repeated until the error stagnates and we have the gEC of the model. The asymmetry of the fitted model is calculated, of which an intuitive overview is given by Figure 1D. As mentioned, the gEC contains the weights of the bidirectional connections, the infrastructure of the model. The PCI and irreversibility are calculated from the simulated data. As shown in Figure 1E one of the nodes in the model is perturbed by changing the value of the bifurcation parameter for a period. This perturbation influences the rest of the model through its connections, creating a pre- and a postperturbational time series, i.e. a baseline and a response. The irreversibility is calculated on the preperturbational time series.

### gEC asymmetry characterises different states of reduced consciousness

The asymmetry of the personalised generative Effective Connectivity (gEC) matrices was calculated, yielding one value per subject. As can be seen in Figure 2A and 2B the higher levels of consciousness were found to have a higher asymmetry in their networks. To calculate whether the distributions are significantly different from each other we performed a non-parametric permutation test (Wilxocon ranksum test), with a Benjamini-Hochberg correction for multiple comparisons for the DoC dataset, and we tested the effect size using Cohen’s d (20). For the Disorders of Consciousness (DoC) dataset the distribution of the models fitted to the CNT subjects has significantly higher asymmetry values than the distribution of models fitted to the UWS subjects (p = 0.0043, Cohen’s d = 1.34). Between the MCS and UWS subjects (p = 0.0564, Cohen’s d = 0.44) and between the MCS and CNT subjects (p = 0.0820, Cohen’s d = 0.65) no significant differences were found.

**Figure 2.**
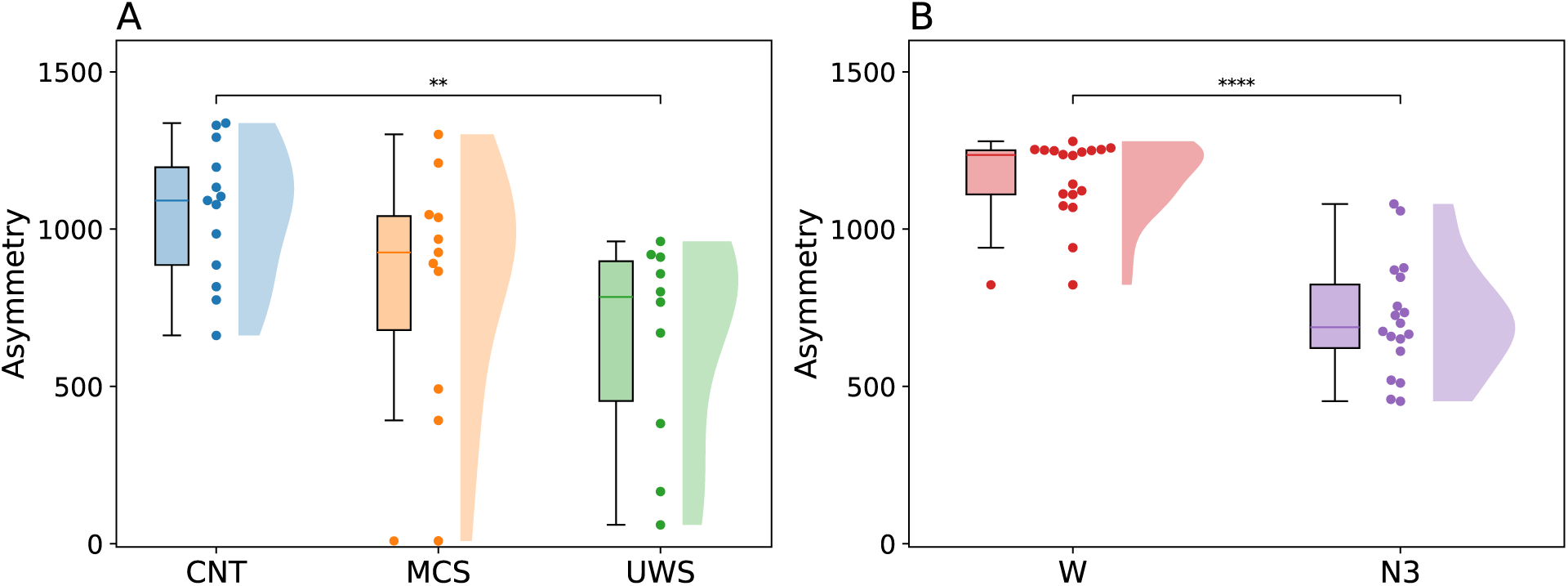
Comparing the level of asymmetry in the gEC for different states of reduced consciousness. Asymmetry is calculated for each gEC matrix and thus gives us one value per subject, per state. A) The asymmetry values for the DoC dataset. B) The asymmetry values for the sleep dataset. CNT - control, MCS - Minimally Conscious State, UWS - Unresponsive Wakefulness State, W – wakefulness, N3 – deep sleep state.

For the sleep dataset the difference between the two states W and N3 was found to be significant (p < 0.001, Cohen’s d = 2.65). These results suggest that models fitted to subjects residing in higher levels of consciousness tend to have higher asymmetry in the gEC. The average asymmetry value of the MCS distribution does not have a significantly different distribution from the CNT and UWS distributions, but the average does fall between the averages of the CNT and UWS distributions, as expected following this line of thought.

### The impact of gEC on the whole-brain nonequilibrium dynamics

As stated before, we used the gEC to inform the connectivity of the model, creating one model per subject (*C*^*ij*^ in Eq. 2 and Eq. 3). To look into the level of nonequilibrium of each model, the irreversibility of the preperturbational time series, i.e. the resting state dynamics, was calculated using the INSIDEOUT framework (18). We ran ten simulations per model for this, yielding 130, 110, 100 and 180 values for subjects in the CNT, MCS, and UWS categories, and the sleep dataset, respectively. As can be seen in Figure 3A all states were significantly different from each other. The difference between CNT and MCS was significant (p = 0.008, Cohen’s d = 0.419), as were the differences between MCS and UWS distributions (p = 0.002, Cohen’s d = 0.412), and between the CNT and UWS distributions (p < 0.001, Cohen’s d = 0.704). These results indicate the tendency of irreversibility diminishing with lower levels of consciousness, as has been previously shown (19), indicating a state closer to equilibrium. The difference between the irreversibility values of W and N3 were significant as well, where the lower level of consciousness again had a lower level of irreversibility (p < 0.001, Cohen’s d = 0.799), which can be observed in Figure 3B.

**Figure 3.**
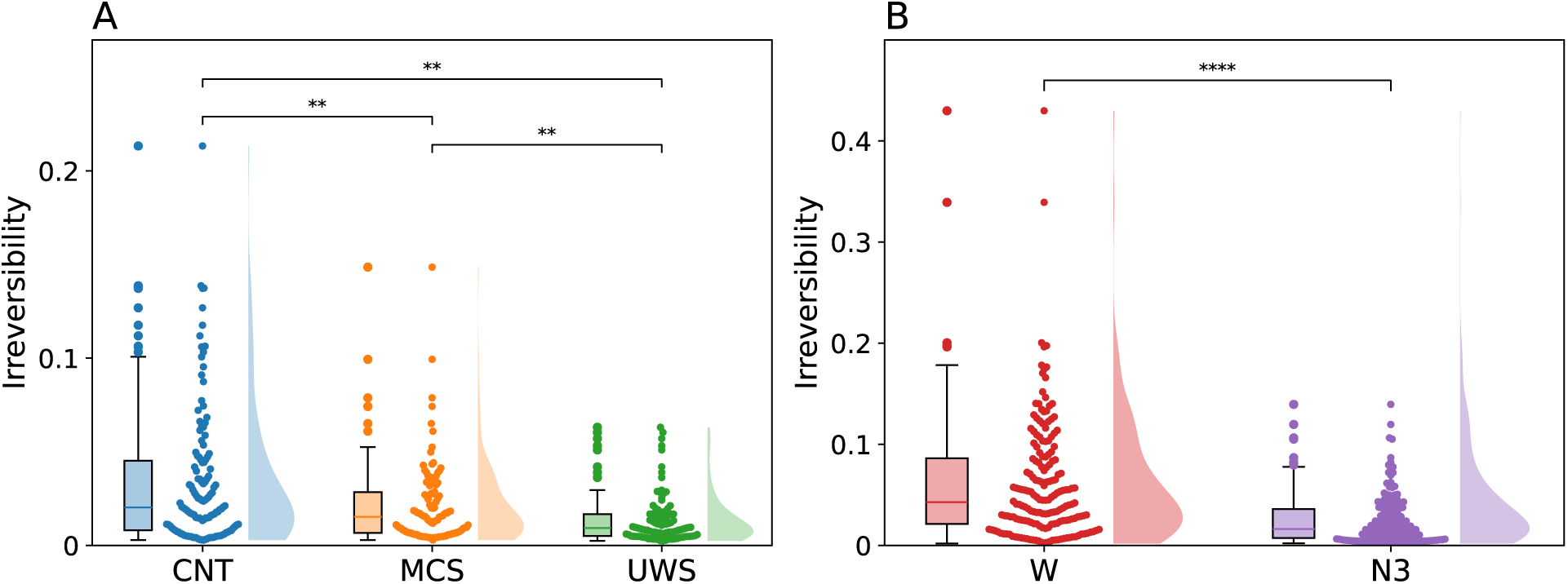
The underlying gEC generates different whole brain nonequilibrium dynamics. The INSIDEOUT framework is applied to the simulated time series of each model to calculate the irreversibility. Ten simulations are performed per model. A) The irreversibility values for the DoC dataset. B) The irreversibility values for the sleep dataset. CNT - control, MCS - Minimally Conscious State, UWS - Unresponsive Wakefulness State, W – wakefulness, N3 – deep sleep state

### Perturbative Complexity

To calculate the PCI values for the simulations both the preperturbational and the postperturbational time series were used to calculate the State Transition PCI. A simulation was performed for each node for each of the models, which are depicted individually in the boxplots shown in Figure 4, meaning there are 90 PCI values depicted for each subject.

**Figure 4.**
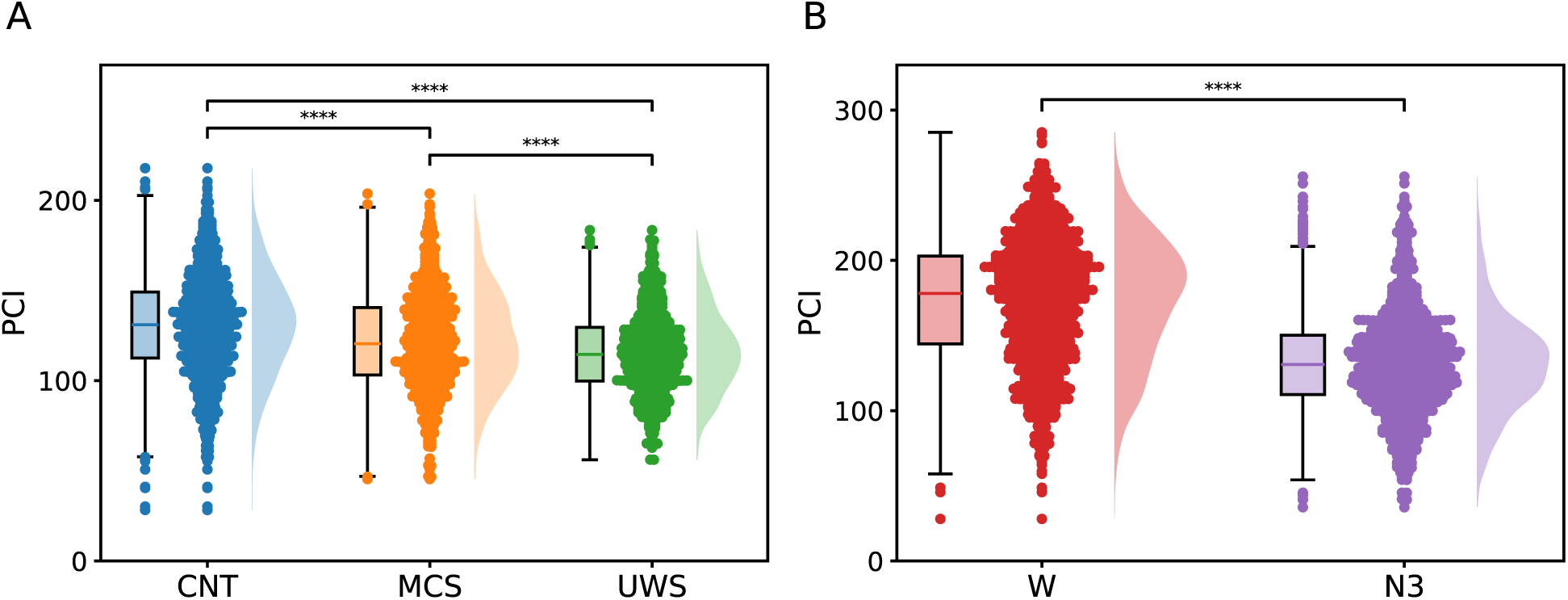
The underlying gEC generates different PCI values in different levels of consciousness. The PCI was calculated on simulated perturbations in the model. One node was perturbed per simulation. The figure shows a PCI value for each node and each subject. A) The PCI values for the DoC dataset. B) The PCI values for the sleep dataset. CNT - control, MCS - Minimally Conscious State, UWS - Unresponsive Wakefulness State, W – wakefulness, N3 – deep sleep state

Significant differences were found in all three comparisons, after Benjamini-Hochberg correction for multiple comparisons, namely the comparison between the MCS and the UWS group (p < 0.001, Cohen’s d = 0.250), the comparison between CNT and MCS (p < 0.001, Cohen’s d = 0.307), and the comparison between CNT and UWS (p < 0.001, Cohen’s d = 0.570). Similar to the irreversibility and asymmetry values the PCI has a tendency to be higher in models that were fitted to subjects residing in higher levels of consciousness. This can be confirmed by looking at Figure 4B where this trend is more pronounced, where the values for the W and N3 state are portrayed (p < 0.001, Cohen’s d = 1.129).

### Relation between asymmetry, irreversibility, and PCI

We performed a hundred simulations of perturbations per node per model to calculate the PCI. We then averaged the PCI values per model, i.e. we have one PCI value per model. To calculate the irreversibility, we performed a hundred simulations per model and calculated the average, yielding one irreversibility and one PCI value per model. We calculated the Spearman’s correlation between the asymmetry and irreversibility, the PCI and irreversibility, and the PCI and the asymmetry. In Figure 5 and 6 the scatterplots are shown together with their Spearman’s correlation values, and the p-value based on a permutation test.

**Figure 5.**
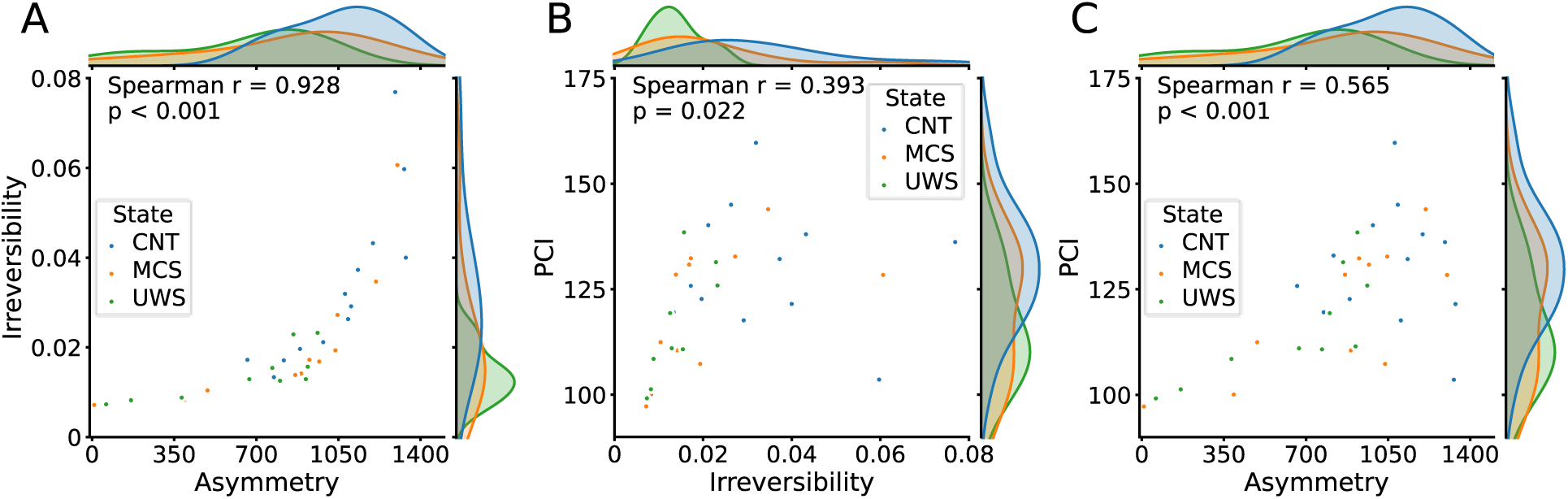
Relations between the irreversibility, asymmetry, and PCI for disorders of consciousness expressed by Spearman’s correlation. Spearman’s correlations between the asymmetry, irreversibility, and PCI for the simulations from the models fitted to the subjects in the DoC dataset. Hundred simulations were averaged over all ROI’s per subject, where the PCI values as well as the irreversibility values were averaged per subject. A) The average PCI per subject is set out against the corresponding average irreversibility. B) The asymmetry of the gEC is set out against the corresponding average irreversibility. C) The asymmetry of the gEC is set out against the corresponding average PCI values. CNT - control, MCS - Minimally Conscious State, UWS - Unresponsive Wakefulness State.

Figures 5 and 6 show that there appears to be a relation between the irreversibility and the asymmetry. This is reflected in the particularly strong correlation values, which were found to be significant in both datasets. For the DoC dataset, this relation is captured by a high correlation (Spearman r = 0.928, p < 0.001), and can similarly be found in the sleep dataset (Spearman r = 0.868, p < 0.001). These correlations across datasets suggest that irreversibility scales with asymmetry, regardless of the type of reduced consciousness. The relation between PCI and irreversibility seems to be a bit more variable, but found to be significant in both datasets after Benjamini-Hochberg correction for multiple comparisons. In the DoC dataset the correlation is weaker (Spearman r = 0.393, p = 0.022), whereas in the sleep dataset the correlation is stronger (Spearman r = 0.686, p < 0.001).

**Figure 6.**
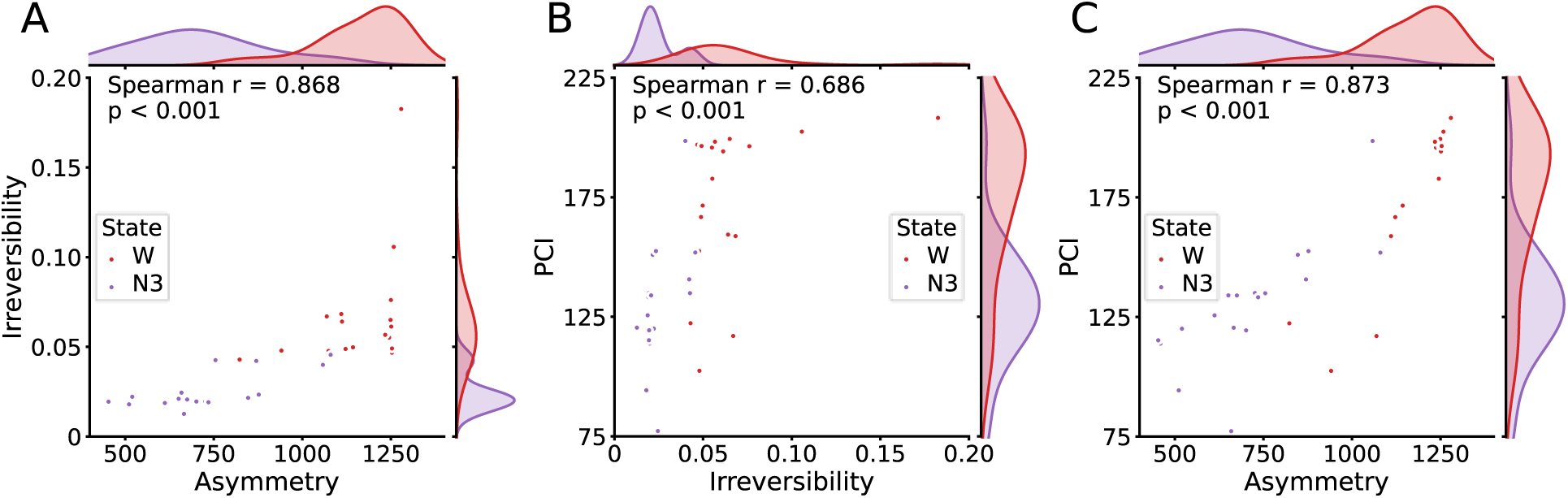
Relations between the irreversibility, asymmetry and PCI for wakefulness and deep sleep expressed by Spearman’s correlation. Spearman’s correlations between the asymmetry, irreversibility, and PCI, for the simulations from the models fitted to the subjects of the sleep dataset. Hundred simulations were averaged over all ROI’s per subject, where the PCI values as well as the irreversibility values were averaged per subject. A) The average PCI per subject is set out against the corresponding average irreversibility. B) The asymmetry of the gEC is set out against the corresponding average irreversibility. C) The asymmetry of the gEC is set out against the corresponding average PCI values. W - wakefulness, N3 - deep sleep state.

Lastly, PCI and asymmetry are found to have a positive significant correlation in both datasets. For the DoC dataset, this correlation is slightly lower (Spearman r = 0.565, p < 0.001) than the strong correlation in the sleep dataset (Spearman r = 0.873, p < 0.001). This indicates that across states of consciousness, higher asymmetry is associated with greater complexity in response to stimuli.

Taken together, our results indicate that higher asymmetry is associated with higher values of PCI and irreversibility. This leads us to state that there is a strong and significant correlation between these measures that was not previously reported on in this setup.

Linear mixed effect (LME) models were created and compared to check whether the states of the subjects influence the correlation between the asymmetry and the other metrics. The LME models with the states taken as a random effect were not significantly different from the LME models without a random effect, as per a likelihood ratio test (LRT), meaning the correlations shown do not appear to be influenced by the states of the subjects.

## Discussion

Our computational modelling approach illuminates how the human brain complexity induced by external stimuli depend on the nonequilibrium dynamics of the brain before being perturbed. Leveraging the thermodynamical description of nonequilibrium in brain dynamics (12,22) and computational whole-brain modelling (23,24), we quantified how the level of nonequilibrium of brain dynamics is related to different states of reduced consciousness and, in turn, how these differences shape the brain’s capability to respond to an external stimulus measured by the PCI.

Our goals were twofold. First, we sought to determine whether the level of asymmetry captured in the inferred generative connectivity between brain regions (20) can represent a signature of consciousness, and whether this asymmetry determines the level of nonequilibrium on brain signals. Second, we aimed to determine whether the generative effective connectivity (gEC) is also driving the perturbational complexity index of consciousness (PCI), establishing a bridge between the nonequilibrium nature of the unperturbed system and the elicited complexity after the perturbation.

Thanks to the recent developments in computational modelling and fMRI data acquisition, we were able to characterise the individual brain dynamics of participants with different states of reduced consciousness by fitting their gEC. Importantly, the asymmetry in the gEC is known to break the detailed balance, driving the system out of equilibrium (14,20). We showed here that higher levels of asymmetry are related to higher levels of consciousness. Importantly, we observed that the level of asymmetry decreases both for DoC patients and for participants in deep sleep, indicating that such changes are not specific to a particular reduced state of consciousness but are associated with the level of consciousness itself. Previous work has consistently shown the relation between nonequilibrium brain dynamics and reduced consciousness using empirical data (13,18,19) as well as through whole-brain models (14,25).

It has been widely investigated how the dynamics of the human brain are constrained and supported by the structural connectome (26–29). In this sense, whole-brain models provided a suitable and successful avenue to investigate how the function of the brain is shaped by its structure (23,27,30,31). Following the same rationale, we used the generative capabilities of whole-brain models to investigate how the gEC shapes the nonequilibrium of the brain dynamics and how that changes between states of consciousness. Crucially, we found a positive correlation between gEC asymmetry and the level of nonequilibrium in brain dynamics at subject level, suggesting that the broken detailed balance in the generative space determines the level of nonequilibrium at the whole-brain level, as has been demonstrated in other biological systems (32). Our results also showed that the level of nonequilibrium in brain dynamics obtained with different gEC’s significantly decreases as the state of consciousness decreases, aligning with empirical results.

In particular, within the thermodynamics description of brain dynamics, FDT establishes a direct link between the nonequilibrium nature of the system and its response to an external stimulus (14,15,25,33). This approach provides a necessary theoretical framework for the very influential papers on PCI by Massimini and colleagues. They empirically demonstrated that the complexity after perturbations can be used as biomarkers of consciousness using transcranial magnetic stimulation (TMS) and electroencephalography (EEG) (4–6). In their work, Massimini and colleagues defined a perturbational complexity index (PCI) that directly measures the amount of information contained in the perturbation-evoked responses by calculating the Lempel-Ziv complexity in space and time of the EEG signals (4). Following the same principals, they addressed the limitations of the Lempel Ziv PCI and defined the State Transition PCI, using dimensionality reduction and state transition quantification (34). These PCI measures have been successfully used for separation of brain states in healthy subjects during wakefulness, dreaming, sleep, under different levels of anaesthesia, and in comatose patients (4–6). Here, we leveraged whole-brain models to compute the State Transition PCI by systematically and perturb our brain models *in silico*. While previous works computed the PCI at the group level (35–38), crucially, here we computed an individualized PCI based on the gEC obtained for each participant in each condition. We found that our computational PCI confirms the empirical results obtained in previous works (4). Importantly, as our computational approach allows us to systematically perturb all regions in the brain, we found that the PCI decreases as the state of consciousness decreases, independently of the perturbed brain region.

The primary benefit of our proposal compared to earlier empirical methods is that it offers insights into the causal generative mechanisms of complex brain dynamics across various brain states. Crucially, we found that the asymmetry of the gEC underlying the brain dynamics highly correlates with the PCI across DoC patients and participants falling asleep, regardless of the perturbed node. This is of particular relevance because, as mentioned in Virmani et al. 2018, an exhaustive manner to perturb the brain of patients is not achievable with the TMS-EEG setup. In this study we show a feasible alternative by using an *in silico* approach (39). They also state that the PCI is dependent on the current state of the system, in line with our conclusion that the PCI is a product of the state of the system, with the notion that the state of the system is defined by the level of nonequilibrium. In a recent paper of Casarotto et al. 2024 the authors reflect on some shortcomings of the PCI, mentioning that it still needs trained experts to perform this test, and not all patients are eligible for TMS procedures, limiting its usefulness (40). They also note several strict criteria to which this TMS-EEG need to be upheld. Following the line of thought that the PCI is determined by the level of nonequilibrium, indicates that the TMS-EEG setup could be redundant.

The link between the empirically performed perturbations and PCI values and their simulated equivalents is beyond the scope of this work and should be investigated in future research. Previous studies using empirical data have consistently shown that decreased levels of consciousness are associated with lower PCI values and reduced irreversibility levels (8,19,34). This leads us to the hypothesis that there is a strong correlation between the empirical PCI and the metrics presented here. Importantly, future work should focus on investigating this relationship based on empirical measures of non-equilibrium dynamics and FDT approaches (25,41) with the empirical PCI.

Taken together, these findings indicate that the broken detailed balance of the brain dynamics, reflected in the asymmetry of the underlying gEC, can be used as a model-based biomarker of consciousness. In turn, this asymmetry lies at the root of the difference in nonequilibrium and PCI, establishing a direct link between the nonequilibrium of the unperturbed system and its responsiveness to external stimuli. Additionally, it could reduce the need for costly experiments, enhance statistical reliability, and minimize potential ethical issues.

## Materials and methods

### Disorders of Consciousness data

This research was approved by the local ethics committee Comité de Protection des Personnes Ile de France 1 (Paris, France) under the code ‘Recherche en soins courants’ (NEURODOC protocol, no. 2013-A01385-40). The patients’ relatives gave their informed consent for their familiar to participate, and all investigations were performed according to the Declaration of Helsinki and the French regulations.

We used fMRI data from 21 patiens with DoC’s. 10 of those patients were diagnosed with UWS and the other 11 patients were diagnosed with MCS. fMRI data was collected from an additional 13 healthy control subjects. This dataset comes from a larger dataset previously described in Escrichs et al. 2022 (42). Trained clinicians conducted the clinical assessment and CRS-R scoring to determine the patients’ level of consciousness. Patients were diagnosed with MCS if they showed some behaviour that could be indicative of awareness, such as visual pursuit, orientation to pain, or reproducible command following. Patients were diagnosed with UWS if they showed signs of arousal (through opening their eyes) without any signs of awareness (never showing non-reflex voluntary movements).

The fMRI data for this dataset were acquired with a 3T General Electric Signa System. T2*-weighted whole-brain resting state images were captured with a gradient-echo EPI sequence using axial orientation (200 volumes, 48 slices, slice thickness: 3 mm, TR/TE: 2400 ms/30 ms, voxel size: 3.4375 × 3.4375 × 3.4375 mm, flip angle: 90°, FOV: 220 mm^2^). An anatomical volume was obtained using a T1-weighted MPRAGE sequence in the same acquisition session (154 slices, slice thickness: 1.2 mm, TR/TE: 7.112 ms/3.084 ms, voxel size: 1 × 1 × 1 mm, flip angle: 15°).

Pre-processing of the resting state data was performed using FSL (http://fsl.fmrib.ox.ac.uk/fsl) as described previously (42). Resting state fMRI was computed using MELODIC (multivariate exploratory linear optimized decomposition into independent components) (43). Steps included discarding the first five volumes, motion correction using MCFLIRT (44), brain extraction tool (BET) (45), spatial smoothing with 5 mm FWHM Gaussian kernel, rigid-body registration, high pass filter cutoff at 100.0 s, and single-session independent component analysis (ICA) with automatic dimensionality estimation. Lesion-driven artefacts (for patients) and noise components were regressed out independently for each subject using FIX (FMRIB’s ICA-based X-noiseifier) (46). The time series were extracted in the AAL parcellation (21).

### Sleep and wakefulness data

Written informed consent was obtained, and the study was approved by the ethics committee of the Faculty of Medicine at the Goethe University of Frankfurt, Germany. We used fMRI data from 18 subjects residing in both wakefulness (W) and deep sleep (N3), previously described in (14). The data was acquired using a 3T system (Siemens Trio, Erlangen, Germany) with settings: 1505 volumes of T2*-weighted echo planar images with a repetition time (TR) of 2.08 seconds, and an echo time of 30 ms; matrix 64 x 64, voxel size 3 x 3 x 2 mm^3^, distance factor 50%, field of view (FOV) 192 mm2.

The EPI data were realigned, normalised to MNI space, and spatially smoothed using a Gaussian kernel of 8 mm^3^ FWHM in SPM8 (http://www.fil.ion.ucl.ac.uk/spm/). Spatial downsampling was then performed to a 4 x 4 x 4 mm^3^ resolution. From the simultaneously recorded ECG and respiration, cardiac- and respiratory-induced noise components were estimated using the RETROICOR method (47), which were regressed out of the signals together with motion parameters. The data were temporally band-pass filtered in the range 0.008-0.08 Hz using a sixth-order Butterworth filter. The time series were extracted in the AAL parcellation (21).

Simultaneous polysomnography (PSG) was performed by recording EEG, EMG, ECG, EOG, pulse oximetry, and respiration. EEG was recorded using a cap (modified BrainCapMR, Easycap, Herrsching, Germany) with 30 channels, where the FCz electrode was used as reference. The sampling rate of the EEG was 5 kHz, and a low-pass filter was applied at 250 Hz. MRI and pulse artefact correction were applied based on the average artefact subtraction method (24) in Vision Analyzer2 (Brain Products, Germany). EMG was collected with chin and tibial derivations, while the ECG and EOG rec orded bipolarly at a sampling rate of 5 kHz with a low-pass filter at 1 kHz. Pulse oximetry was collected using the Trio scanner, and respiration was recorded with MR-compatible devices (BrainAmp MR+, BrainAmp ExG; Brain Products, Gilching, Germany).

Participants were instructed to lie still in the scanner with their eyes closed and relax. Sleep classification was performed by a sleep expert based on the EEG recordings in accordance with the AASM criteria (2007).

### Structural connectivity

We used the SC of the 90 ROI’s as presented in a previous study, for a more detailed description please refer to (48). Briefly, a SC matrix was obtained for each subject (n = 16) by using tractography algorithms to Diffusion Tensor Imaging, following a previously described methodology (49). The connection between regions C_ij_ was calculated as the proportion of sampled fibres in all voxels in region i that reach any voxel in region j. DTI does not portray directionality of these fibres, therefore the average was taken between C_ij_ and C_ji_ as the undirected connection between regions i and j. The average was taken over the 16 subjects to portray a SC matrix representing a healthy connectivity.

### Statistical testing and multiple comparison correction

To test whether two distributions are significantly distinct from one another we used a permutation based ranksum t-test with 10,000 permutations. To correct for multiple comparisons, we used a Benjamini-Hochberg procedure, where the significance threshold is adjusted for each p-value based on their rank in order of magnitude. The threshold is then determined by dividing the p-value by the number of tests performed and multiplying by its rank.

To test whether two metrics are correlated we used Spearman’s correlation to check for monotonic correlations and a non-parametric permutation-based test with 10,000 permutations. Again, significance threshold for the p-values were corrected through a Benjamini-Hochberg procedure.

### Effect size

To calculate and interpret the effect size of the difference between two distributions we used Cohen’s d effect size (50). We adjusted the effect sizes where needed when dealing with smaller sample sizes (N < 50) to correct for possible instability in small sample sizes.

### Whole-brain modelling

The simulations are done by using the Hopf model, a model of coupled Stuart Landau oscillators. These oscillators are described by the normal form of a supercritical Hopf bifurcation that represent the different ROI’s. The ROIs, i.e. the nodes in the model, are connected through a Structural Connectivity (SC) derived from empirical data. The bifurcation parameter of the Hopf bifurcation is set to a value close to 0 so that the oscillator portrays the oscillatory and noisy behaviour that is characteristic for the brain (23). In Eq. 1 the dynamics of one oscillator are depicted in its uncoupled form, where z_j_ represents the dynamics as a complex value (with z_j_ x_j_ + y_j_). In this equation *ω* represents the intrinsic frequency of the node, calculated from the fMRI data as the averaged peak frequency and bandpass filtered between 0.04 and 0.07 Hz, and *a* the bifurcation parameter which was set at -0.02:<colcnt=1>

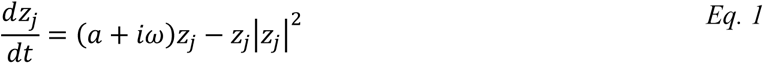

As previously mentioned, the oscillators are connected through an infrastructure represented by an SC, depicted as *C*_*ij*_ in Eq. 2 and Eq. 3:

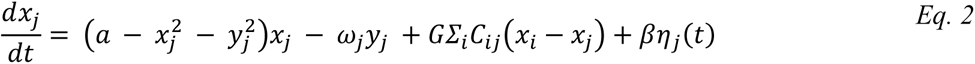

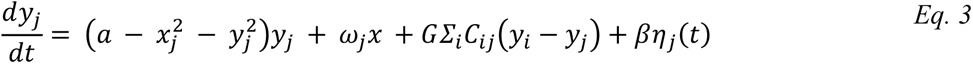

Similar as per Eq. 1 *a* represents the bifurcation parameter and ω represents the intrinsic frequency of the node, and η noise. The global coupling factor is represented by G and in this work it is kept at a value of 1 for all models. For a more detailed description of the Hopf model, please refer to Deco et al. 2017 (23).

### Generative Effective Connectivity

Each subject’s data was used to fit a generative Effective Connectivity (gEC) to create personalised models by using the linear approximation of the Hopf model (51), depicted in Figure 1C. All data were band-pass filtered between 0.04-0.07 Hz before fitting. A model is fitted to the empirical time-lagged covariances, depicted in Eq. 6, by iteratively adjusting the values of the connections in C_ij_ which are used in Eq. 2 and Eq. 3 to simulate time series. The personalisation of the models was done by first creating grouped models. The time-lagged covariance matrix of each subject was calculated and averaged per group, yielding five grouped models: three grouped models for the DoC dataset (CNT, MCS, UWS) and two grouped models for the sleep dataset (W, N3). From there on each personalised model was created by taking the gEC of their respective grouped model as an initial step for the fitting process. An SC mask is used, meaning that all gECs in the end have connections in the same constellation but not with the same weights.

### Asymmetry

The asymmetry of a model was calculated by looking at the number of pairs that were asymmetrically connected in the gEC. An intuitive overview of this metric is shown in Figure 1D. The transposed gEC was subtracted from its original form, yielding a symmetric matrix. The absolute values of this symmetric matrix were binarized by using a threshold, 0 for values below the threshold and 1 for values above the threshold, as per Eq. 4. The asymmetry value depicts the number of pairs of nodes, and thus the sum of the lower triangle was taken as the number of pairs that were connected sufficiently asymmetric to survive the thresholding. Mathematically this can be seen as summing matrix A yielded in Eq. 4 and divided by two as per Eq. 5.

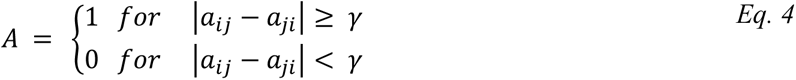

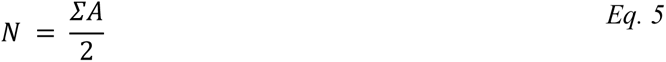

The threshold was generalized for all models and set so that all asymmetry values were at least non-zero. In other words, all gECs should have at least one pair of nodes that was connected asymmetrically enough to surpass the threshold, which was found to be at a value of 0.12.

### Insideout

To compute the irreversibility of the resting state dynamics of the models we used the INSIDEOUT framework (18). This framework looks at the temporal asymmetry of time series by comparing the time shifted covariance matrices of the forward and the reversed time series, as shown in Eq. 6:

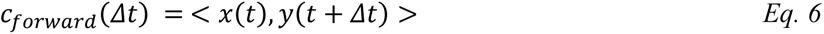

By looking at the distance of these covariance matrices, a value is calculated to portray how high the level of irreversibility is in the system, this depicts the arrow of time of the system and represents the level of non-equilibrium. For a multidimensional system this equation looks like:

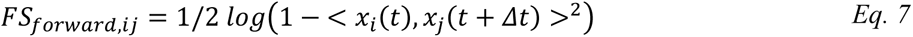

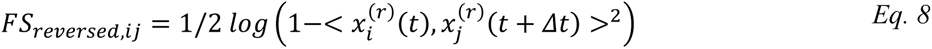

Where the irreversibility is calculated as the mean of the absolute squares of the elements of the difference between FS_forward_ and FS_reversed_:

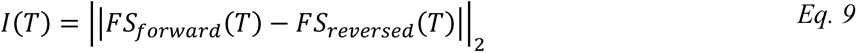

For a more detailed description of this framework, please refer to Deco et al. 2022 (18).

### PCI simulated

To introduce perturbations in the model we made use of the Hopf model’s bifurcation characteristics. By changing the value of the bifurcation parameter to a positive value a node is pushed into an oscillatory state. By doing so, the oscillatory activity of this node will affect the other nodes based on their connectedness as dictated by the gEC. The preperturbational time series is used as a baseline and the postperturbational time series is used as a response window as defined by the methodology of the State Transition PCI. A principal component analysis is done on the response window and by using a singular value decomposition the baseline is expressed in these principal components as well. The principal components that make up 99% of the response are the ones that are used in the analysis of the number of state transitions. The PCI is the summed difference in number of state transitions between the baseline and the response window expressed in principal components. For a more detailed description of the State Transition PCI please refer to Comolatti et al. 2019 (34).

## Data and code availability

The code to simulate the model including the perturbations, and calculate the measures presented in this paper are available at https://github.com/wiepstikvoort/Nonequilibrium-asymmetry-PCI

The disorder of consciousness datasets contain information from a clinical population and are not publicly available due to constraints imposed by the approved ethics protocol.

The sleep data set is publicly available at https://github.com/yonisanzperl/Perturbation_in_dynamical_models

## Acknowledgements

**W.S.** is an FI fellow with the support of AGAUR, Generalitat de Catalunya and Fondo Social Europeo (2022 FI_B 00152). **E.P.** is an FPI fellow funded by the Spanish "Ministerio de Ciencia, Innovación y Universidades" (MICIU/AEI/10.13039/501100011033) and "ESF investing in your future" under the grant PRE2020-0961. **I.M.** is funded by FLAG-ERA research funding organisation (project ModelDXConsciousness). **G.D. and A.E.** were supported by the Grant PID2022-136216NB-I00 funded by MICIU/AEI/10.13039/501100011033 and by “ERDF A way of making Europe,” ERDF, EU. **G.D. and Y.S.P.** were supported by the project NEurological MEchanismS of Injury, and Sleep-like cellular dynamics (NEMESIS) (ref. 101071900) funded by the EU ERC Synergy Horizon Europe. **A.E.** was also supported by the project eBRAIN-Health - Actionable Multilevel Health Data (id 101058516), funded by the EU Horizon Europe. **Y.S.P.** is also supported by the European Union’s Horizon 2020 research and innovation program under the Marie Sklodowska-Curie grant 896354. **G.D.** is also supported by AGAUR research support grant (2021 SGR 00917) funded by the Department of Research and Universities of the Generalitat of Catalunya. **M.L.K.** is supported by the Centre for Eudaimonia and Human Flourishing (funded by the Pettit and Carlsberg Foundations) and Center for Music in the Brain (funded by the Danish National Research Foundation, DNRF117). **J.D.S.** is supported by the EU ERA PerMed Joint Translational 2019 project (project PerBrain) and by the JTC-HBP project MODELDxConsciousness.

## Notes

https://github.com/wiepstikvoort/Nonequilibrium-asymmetry-PCI

### Competing Interest Statement

The authors have declared no competing interest.

## References

1. Sarasso S, Casali AG, Casarotto S, Rosanova M, Sinigaglia C, Massimini M. Consciousness and complexity: a consilience of evidence. Neurosci Conscious. 2021 Aug 30;niab023.

2. Deco G, Tononi G, Boly M, Kringelbach ML. Rethinking segregation and integration: contributions of whole-brain modelling. Nat Rev Neurosci. 2015 Jul;16(7):430–9.

3. Chialvo DR. Emergent complex neural dynamics. Nat Phys. 2010 Oct;6(10):744–50.

4. Casali AG, Gosseries O, Rosanova M, Boly M, Sarasso S, Casali KR, et al. A Theoretically Based Index of Consciousness Independent of Sensory Processing and Behavior. Sci Transl Med. 2013 Aug 14;5(198).

5. Ferrarelli F, Massimini M, Sarasso S, Casali A, Riedner BA, Angelini G, et al. Breakdown in cortical effective connectivity during midazolam-induced loss of consciousness. Proc Natl Acad Sci. 2010 Feb 9;107(6):2681–6.

6. Massimini M, Ferrarelli F, Huber R, Esser SK, Singh H, Tononi G. Breakdown of Cortical Effective Connectivity During Sleep. Science. 2005 Sep 30;309(5744):2228–32.

7. Sarasso S, Rosanova M, Casali AG, Casarotto S, Fecchio M, Boly M, et al. Quantifying Cortical EEG Responses to TMS in (Un)consciousness. Clin EEG Neurosci. 2014 Jan;45(1):40–9.

8. Sinitsyn DO, Poydasheva AG, Bakulin IS, Legostaeva LA, Iazeva EG, Sergeev DV, et al. Detecting the Potential for Consciousness in Unresponsive Patients Using the Perturbational Complexity Index. Brain Sci. 2020 Nov 27;10(12):917.

9. Wang Y, Niu Z, Xia X, Bai Y, Liang Z, He J, et al. Application of Fast Perturbational Complexity Index to the Diagnosis and Prognosis for Disorders of Consciousness. IEEE Trans Neural Syst Rehabil Eng. 2022;30:509–18.

10. Casarotto S, Comanducci A, Rosanova M, Sarasso S, Fecchio M, Napolitani M, et al. Stratification of unresponsive patients by an independently validated index of brain complexity. Ann Neurol. 2016 Nov;80(5):718–29.

11. Lynn CW, Cornblath EJ, Papadopoulos L, Bertolero MA, Bassett DS. Broken detailed balance and entropy production in the human brain. Proc Natl Acad Sci. 2021 Nov 23;118(47):e2109889118.

12. Kringelbach ML, Sanz Perl Y, Deco G. The Thermodynamics of Mind. Trends Cogn Sci. 2024 Jun;28(6):568–81.

13. Sanz Perl Y, Bocaccio H, Pallavicini C, Pérez-Ipiña I, Laureys S, Laufs H, et al. Nonequilibrium brain dynamics as a signature of consciousness. Phys Rev E. 2021 Jul 28;104(1):014411.

14. Deco G, Lynn CW, Sanz Perl Y, Kringelbach ML. Violations of the fluctuation-dissipation theorem reveal distinct nonequilibrium dynamics of brain states. Phys Rev E. 2023 Dec 26;108(6):064410.

15. Crisanti A, Ritort F. Violation of the fluctuation-dissipation theorem in glassy systems: basic notions and the numerical evidence. J Phys Math Gen. 2003 May 30;36(21):R181–290.

16. Onsager L. Reciprocal Relations in Irreversible Processes. I. Phys Rev. 1931 Feb 15;37(4):405–26.

17. Onsager L. Reciprocal Relations in Irreversible Processes. II. Phys Rev. 1931 Dec 15;38(12):2265–79.

18. Deco G, Sanz Perl Y, Bocaccio H, Tagliazucchi E, Kringelbach ML. The INSIDEOUT framework provides precise signatures of the balance of intrinsic and extrinsic dynamics in brain states. Commun Biol. 2022 Jun 10;5(1):572.

19. G-Guzmán E, Perl YS, Vohryzek J, Escrichs A, Manasova D, Türker B, et al. The lack of temporal brain dynamics asymmetry as a signature of impaired consciousness states. Interface Focus. 2023 Jun 6;13(3):20220086.

20. Kringelbach ML, Perl YS, Tagliazucchi E, Deco G. Toward naturalistic neuroscience: Mechanisms underlying the flattening of brain hierarchy in movie-watching compared to rest and task. Sci Adv. 2023 Jan 13;9(2):eade6049.

21. Tzourio-Mazoyer N, Landeau B, Papathanassiou D, Crivello F, Etard O, Delcroix N, et al. Automated Anatomical Labeling of Activations in SPM Using a Macroscopic Anatomical Parcellation of the MNI MRI Single-Subject Brain. NeuroImage. 2002 Jan;15(1):273–89.

22. Lynn CW, Bassett DS. The physics of brain network structure, function and control. Nat Rev Phys. 2019 Mar 27;1(5):318–32.

23. Deco G, Kringelbach ML, Jirsa VK, Ritter P. The dynamics of resting fluctuations in the brain: metastability and its dynamical cortical core. Sci Rep. 2017 Jun 8;7(1):3095.

24. Breakspear M. Dynamic models of large-scale brain activity. Nat Neurosci. 2017 Mar;20(3):340–52.

25. Monti JM, Perl YS, Tagliazucchi E, Kringelbach M, Deco G. The fluctuation-dissipation theorem and the discovery of distinctive off-equilibrium signatures of brain states [Internet]. 2024 [cited 2024 Sep 13]. Available from: http://biorxiv.org/lookup/doi/10.1101/2024.04.04.588056

26. Suárez LE, Markello RD, Betzel RF, Misic B. Linking Structure and Function in Macroscale Brain Networks. Trends Cogn Sci. 2020 Apr;24(4):302–15.

27. Honey CJ, Kötter R, Breakspear M, Sporns O. Network structure of cerebral cortex shapes functional connectivity on multiple time scales. Proc Natl Acad Sci. 2007 Jun 12;104(24):10240–5.

28. Honey CJ, Sporns O, Cammoun L, Gigandet X, Thiran JP, Meuli R, et al. Predicting human resting-state functional connectivity from structural connectivity. Proc Natl Acad Sci. 2009 Feb 10;106(6):2035–40.

29. Luppi AI, Singleton SP, Hansen JY, Jamison KW, Bzdok D, Kuceyeski A, et al. Contributions of network structure, chemoarchitecture and diagnostic categories to transitions between cognitive topographies. Nat Biomed Eng [Internet]. 2024 Aug 5 [cited 2024 Sep 13]; Available from: https://www.nature.com/articles/s41551-024-01242-2

30. Cabral J, Kringelbach ML, Deco G. Functional connectivity dynamically evolves on multiple time-scales over a static structural connectome: Models and mechanisms. NeuroImage. 2017 Oct;160:84– 96.

31. Deco G, Ponce-Alvarez A, Hagmann P, Romani GL, Mantini D, Corbetta M. How Local Excitation-Inhibition Ratio Impacts the Whole Brain Dynamics. J Neurosci. 2014 Jun 4;34(23):7886–98.

32. Battle C, Broedersz CP, Fakhri N, Geyer VF, Howard J, Schmidt CF, et al. Broken detailed balance at mesoscopic scales in active biological systems. Science. 2016 Apr 29;352(6285):604–7.

33. Lindner B. Fluctuation-Dissipation Relations for Spiking Neurons. Phys Rev Lett. 2022 Oct 31;129(19):198101.

34. Comolatti R, Pigorini A, Casarotto S, Fecchio M, Faria G, Sarasso S, et al. A fast and general method to empirically estimate the complexity of brain responses to transcranial and intracranial stimulations. Brain Stimulat. 2019 Sep;12(5):1280–9.

35. Deco G, Cabral J, Saenger VM, Boly M, Tagliazucchi E, Laufs H, et al. Perturbation of whole-brain dynamics in silico reveals mechanistic differences between brain states. NeuroImage. 2018 Apr;169:46–56.

36. Goldman JS, Kusch L, Yalcinkaya BH, Depannemaecker D, Nghiem TAE, Jirsa V, et al. Brain-scale emergence of slow-wave synchrony and highly responsive asynchronous states based on biologically realistic population models simulated in The Virtual Brain [Internet]. 2020 [cited 2024 Sep 13]. Available from: http://biorxiv.org/lookup/doi/10.1101/2020.12.28.424574

37. Kunze T, Hunold A, Haueisen J, Jirsa V, Spiegler A. Transcranial direct current stimulation changes resting state functional connectivity: A large-scale brain network modeling study. NeuroImage. 2016 Oct;140:174–87.

38. Sanz Perl Y, Escrichs A, Tagliazucchi E, Kringelbach ML, Deco G. Strength-dependent perturbation of whole-brain model working in different regimes reveals the role of fluctuations in brain dynamics. Daunizeau J, editor. PLOS Comput Biol. 2022 Nov 2;18(11):e1010662.

39. Virmani M, Nagaraj N. A novel perturbation based compression complexity measure for networks. Heliyon. 2019 Feb;5(2):e01181.

40. Casarotto S, Hassan G, Rosanova M, Sarasso S, Derchi C, Trimarchi PD, et al. Dissociations between spontaneous electroencephalographic features and the perturbational complexity index in the minimally conscious state. Eur J Neurosci. 2024 Mar;59(5):934–47.

41. Patow G, Monti J, Acero-Pousa I, Idesis S, Escrichs A, Perl YS, et al. Off-Equilibrium Fluctuation-Dissipation Theorem Paves the Way in Alzheimer’s Disease Research [Internet]. 2024 [cited 2024 Oct 2]. Available from: http://biorxiv.org/lookup/doi/10.1101/2024.09.15.613131

42. Escrichs A, Perl YS, Uribe C, Camara E, Türker B, Pyatigorskaya N, et al. Unifying turbulent dynamics framework distinguishes different brain states. Commun Biol. 2022 Jun 29;5(1):638.

43. Beckmann CF, Smith SM. Probabilistic Independent Component Analysis for Functional Magnetic Resonance Imaging. IEEE Trans Med Imaging. 2004 Feb;23(2):137–52.

44. Jenkinson M, Bannister P, Brady M, Smith S. Improved Optimization for the Robust and Accurate Linear Registration and Motion Correction of Brain Images. NeuroImage. 2002 Oct;17(2):825–41.

45. Smith SM. Fast robust automated brain extraction. Hum Brain Mapp. 2002 Nov;17(3):143–55.

46. Griffanti L, Salimi-Khorshidi G, Beckmann CF, Auerbach EJ, Douaud G, Sexton CE, et al. ICA-based artefact removal and accelerated fMRI acquisition for improved resting state network imaging. NeuroImage. 2014 Jul;95:232–47.

47. Deco G, Kringelbach ML. Great Expectations: Using Whole-Brain Computational Connectomics for Understanding Neuropsychiatric Disorders. Neuron. 2014 Dec;84(5):892–905.

48. Ipiña IP, Kehoe PD, Kringelbach M, Laufs H, Ibañez A, Deco G, et al. Modeling regional changes in dynamic stability during sleep and wakefulness. NeuroImage. 2020 Jul;215:116833.

49. Cabral J, Kringelbach M, Deco G. Functional Graph Alterations in Schizophrenia: A Result from a Global Anatomic Decoupling? Pharmacopsychiatry. 2012 May;45(S 01):S57–64.

50. Sullivan GM, Feinn R. Using Effect Size—or Why the P Value Is Not Enough. J Grad Med Educ. 2012 Sep 1;4(3):279–82.

51. Ponce-Alvarez A, Deco G. The Hopf whole-brain model and its linear approximation. Sci Rep. 2024 Jan 31;14(1):2615.

